# The uptake of avermectins in *Caenorhabditis elegans* is dependent on Intra-Flagellar Transport and other protein trafficking pathways

**DOI:** 10.1101/2021.10.22.465401

**Authors:** Robert A. Brinzer, David J. France, Claire McMaster, Stuart Ruddell, Alan D. Winter, Antony P. Page

## Abstract

Parasitic nematodes are globally important and place a heavy disease burden on infected humans, crops and livestock, while commonly administered anthelmintics used for treatment are being rendered ineffective by increasing levels of resistance. Although the modes of action and resistance mechanisms caused by detoxification and target site insensitivity for these compounds is well documented, the mechanisms for uptake, which can also cause resistance, are still poorly defined. It has recently been shown in the model nematode *Caenorhabditis elegans* that the avermectins or macrocyclic lactones such as ivermectin and moxidectin gain entry though the sensory cilia of the amphid neurons. This study interrogated the molecular mechanisms involved in the uptake of avermectins using a combination of forward genetics and targeted resistance screening approaches along with visualising a BODIPY labelled ivermectin analog and confirmed the importance of intraflagellar transport in this process. This approach also identified the protein trafficking pathways used by the downstream effectors and the components of the ciliary basal body that are required for effector entry into these non-motile structures. Mutations in many of the genes under investigation also resulted in resistance to the unrelated anthelmintic drugs albendazole and levamisole, giving insights into the potential mechanisms of multidrug resistance observed in field isolates of the parasitic nematodes that are a scourge of ruminant livestock. In total 50 novel *C. elegans* anthelmintic survival associated genes were identified in this study, three of which (*daf-6*, *rab-35* and *inx-19*) are associated with broad spectrum cross resistance. When combined with previously known resistance genes, there are now 53 resistance associated genes which are directly involved in amphid, cilia and IFT function.

**Author Summary:** Nematodes represent significant pathogens of man and domestic animals and control relies heavily on limited classes of Anthelminitic drugs. Single and multi-drug resistance is a growing problem however mechanisms of anthelmintic drug resistance and drug uptake by nematodes remain to be clearly elucidated. In *Caenorhabditis elegans* there has been an association between amphid and dye filling defects with resistance to avermectins however the effector and causal mechanisms remain elusive. This study uses a combination of fluorescently labelled anthelmintics and anthelmintic resistance screens to probe the uptake mechanisms for these drugs. The role of the sensory amphids in the uptake of avermectins was confirmed. The avermectins enter the distal segment of the cilia using an effector which is delivered by the UNC-119 and UNC-33/UNC-44 transport systems to the base of the cilia, followed by distal appendage dependent entry and transport along the cilia by the intraflagellar transport pathway. Of the genes investigated, three (*daf-6*, *rab-35* and *inx-19*) were linked to cross resistance against all the anthelmintics tested (Ivermectin, Moxidectin, Albendazole and Levamisole). This study gives further insight into how important classes of anthelmintics enter nematodes and highlights the potential for this process to give rise to anthelmintic resistance.

## Introduction

Parasitic nematodes place a highly significant and heavy disease burden on infected plants and animals causing annual global yield and productivity losses in excess of $100 billion (1, 2) and in addition requires over $20 billion annually to treat with anthelmintics (3). Currently available broad spectrum anthelmintics are from a limited range of chemical families (3) and resistance to one or more classes is becoming widespread in field populations (4) jeopardizing food security and human health. Therefore, until new anthelmintic classes are developed, it is necessary to prolong the efficacy of existing drugs by finding ways to supress resistance.

The avermectins or macrocyclic lactones such as ivermectin and moxidectin are the most commonly administered anthelmintics due to their low cost and high persistent efficacy (5) however the rapid spread of resistance is beginning to render them ineffective (4). Avermectins function by paralysing the central nervous system, which eventually leads to death, through interaction with multiple subunits of the glutamate gated chloride channel primary target, as well as multiple secondary targets, thereby resulting in constitutive activation (6). The target binding specificity of avermectins is determined by saccharide groups on C-13 (eg. ivermectin), methoxime on C-23 (eg. moxidectin) and alkyl groups on C-25 of the lactone ring (6, 7). Since subunit interactions vary between different avermectins, commonly occurring target site insensitivity mutations in one subunit binding site do not necessarily confer cross resistance to other macrocyclic lactones (8). As nematodes have limited capacity for phase I detoxification of macrocyclic lactones (9, 10), resistance relies on increased phase II conjugation and efflux (11), target site insensitivity or reduced drug uptake (12). However, all identified and candidate resistance genes that interact directly with avermectins or their metabolites function downstream of macrocyclic lactone uptake (11–13). The macrocyclic lactones lack the chemical properties that would allow them to spontaneously cross biological membranes (14) meaning that uptake is dependent on the ability of biological systems of the organism to accumulate appropriate concentrations in the target tissues however, the mechanism and associated genes involved in uptake are still unknown or poorly defined.

There is a high degree of conservation in the layout of the central nervous system between nematode species, which consists of around 200-300 neurons, with sensory inputs from sensilla being processed by the nerve ring to output motor neuron mediated responses (15–17). The amphid sensilla function as the primary sensory organ for environmental stimuli (chemical, ion and osmotic gradients, temperature, pheromones and noxious compounds). The sensilla consist of two pairs of 12-13 neurons (12 in *Caenorhabditis elegans*) which have non-motile cilia enriched in G protein-coupled receptors on the dendrites that are exposed to the environment through pores in the cuticle (16, 18–20). Ciliogenesis of sensory cilia utilise assembly pathways that are conserved throughout Eukaryota where a centriole derived basal body anchors to the cell membrane restricting the local diffusion of proteins and lipids and organises microtubules (21). These microtubules are then used for the delivery of lipids and proteins to the growing cilia by intraflagellar transport (IFT) complexes that travel along the microtubules using dynein and kinesin motors (21).

There has been an observed correlation between macrocyclic lactone resistance caused by reduced uptake and defects in amphid morphology in *Caenorhabditis elegans* with several causative genes being associated with dye-filling, chemosensation, osmosensation, dauer formation and mechanosensation defective phenotypes (12, 22, 23). Amphid morphology and dye filling defects have also been noted in field populations of *Haemonchus contortus* that are resistant to avermectins (22, 24). This current study uses a mechanistic approach to investigate cellular processes associated with previously discovered resistance genes, in combination with targeted resistance screens and a BODIPY labelled ivermectin analog in *C. elegans*, to identify the roles played by anterograde and retrograde intraflagellar transport in the ciliary distal segment of the amphid neurons in the uptake of avermectins (ivermectin and moxidectin). Pathways involved in trafficking cilia proteins to and from the ciliary gate of the basal body were also investigated, revealing that the UNC-101 and UNC-119 mediated secretion pathways and the polarisers of axon-dendrite protein sorting UNC-33 and UNC-44 are important components involved in avermectin uptake, whereas the RAB-35 recycling pathway plays a role downstream of uptake. Candidate avermectin resistance genes were also checked for cross resistance to other anthelmintics, revealing the IFT and the SEC-24 secretion pathways as being important for susceptibility to the benzimidazole drug, albendazole, while susceptibility to the imidazothiazole levamisole is not dependent on any of the tested secretion pathways. A whole genome sequencing approach was applied to map candidates from a forward genetic screen for resistance to avermectins, and in combination with a targeted resistance screen, 50 novel anthelmintic-survival associated genes were uncovered in *C. elegans* including: 16 avermectin resistant, 7 avermectin and albendazole resistant, 9 albendazole resistant, 7 levamisole resistant, 1 avermectin and levamisole resistant and 3 genes (*daf-6*, *rab-35* and *inx-19*) which cause broad spectrum cross resistance to all four anthelmintics tested.

## Methods

### Chemicals

Suppliers and catalogue numbers of all reagents used are listed in the Supplementary Methods.

### Nematode strains

Putative orthologs of key basal body genes for which there was no primary literature were chosen using a combination of Protein BLAST (https://blast.ncbi.nlm.nih.gov/Blast.cgi) and the Marrvel (25) and AceView (26) databases.

TM prefixed strains were obtained from the National BioResource Project, Japan while all other strains used were purchased from the *C. elegans* Genetics Centre, USA. All strains were maintained on *Escherichia coli* OP50-1 inoculated Nematode Growth Medium (NGM) plates following standard protocols (http://www.wormbook.org/toc_wormmethods.html). Strains used in this study are listed in the Supplementary Methods.

### Anthelmintic resistance assays

Anthelmintic stock solutions were prepared as follows: 10µM ivermectin stock was made by the serial dilution of a 10mM stock using DMSO as a solvent for both stocks; 10µM moxidectin stock was prepared using the same procedure as ivermectin; 50mM albendazole stock was made by dissolving in DMSO at 31°C with vigorous agitation; 1M levamisole stock was made by dissolving in sterile distilled water. Stock solutions were dispensed into 1ml aliquots and stored at -20°C.

NGM plates containing anthelmintics were produced by adding volumes of anthelmintic stock solution to cooled molten NGM agar (50°C) before mixing and pouring onto 3cm petri dishes. The volume of anthelmintic stock solution added never exceeded 0.3% of the final volume. Anthelmintic plate concentrations used were 10nM ivermectin, 5nM and 10nM moxidectin, 100µM and 150µM albendazole and 0.2mM and 0.8mM levamisole. Plates were inoculated with 50µl OP50-1 24 hours before starting assays.

To determine ivermectin and moxidectin resistance, survival assays were performed by picking 5 L4 worms of the strain to be tested onto each plate with two biological and two technical replicates. Growth and mortality were inspected every 48 hours using a light microscope. A strain was considered resistant if it could produce an F2 generation before the plates desiccated compared to susceptible strains which showed paralysis and growth arrest with the F1 generation failing to reach adulthood. The wild type N2 strain was used as a susceptible negative control and DA1316(*ad1305*; *vu227*; *pk54*) was used as a resistant positive control. The strength of resistance for ivermectin exposure was categorised as weak (+ (w)) if the population only reached F2, moderate (+) if it reached F3 or F4 and strong (++) if growth was visually indistinguishable from NGM plates without anthelmintics. Moxidectin resistance strength was categorised as weak (+ (w)) if the population only reached F2 on 5nM plates, moderate (+) if it reached F3 or F4 on 5nM plates and strong (++) if it reached F3 or F4 on 10nM plates.

Albendazole and levamisole resistance was determined using uncoordinated phenotype assays by picking 5 adult worms of the strain to be tested onto each plate with two biological and two technical replicates. At day 3 and day 6 a random sample of 20 worms per plate were poked on their head with a platinum wire and scored for the ability to reverse backwards (an inability to reverse corresponds to an Unc or uncoordinated phenotype). For mutant strains which innately show an uncoordinated phenotype (Unc), a different scoring criteria was used; with worms being scored as resistant if any muscle movement was shown in response to being poked and scored as sensitive if they were completely paralysed. N2 was used as a negative (sensitive) control and CB3474(*e1880*) (for albendazole resistance), ZZ1(*x1*) or ZZ15(*x15*) (for levamisole resistance) were used as positive controls. Strains were considered resistant if over 50% of sampled worms were unaffected by the anthelmintic; categorised as moderately resistant (+) at the lower dose and strongly resistant (++) at the higher dose. If at the higher dose a strain was 100% unaffected it was classed as extremely resistant (+++). Susceptible strains which had less than 50% of the sampled population unaffected at the lower dose (-) were categorised as highly susceptible (--) by comparing for impaired growth and reproduction relative to controls on NGM plates without anthelmintics. Strains were deemed extremely susceptible (---) if mortality was observed at either anthelmintic concentration.

### Synthesis and evaluation of BODIPY labelled anthelmintic analogs

Details of chemical synthesis, purification and analysis of fluorescent analogs of ivermectin and albendazole (Fig 1) are listed in Supplementary Methods. <u>F</u>atty-<u>B</u>ODIPY-Ivermectin (FBI) was synthesised in 11 steps as shown in Fig S3A. <u>B</u>ODIPY-<u>AlB</u>enda<u>Z</u>ole (BABZ) was synthesised in 5 steps as shown in Fig S3B.

**Fig 1.**
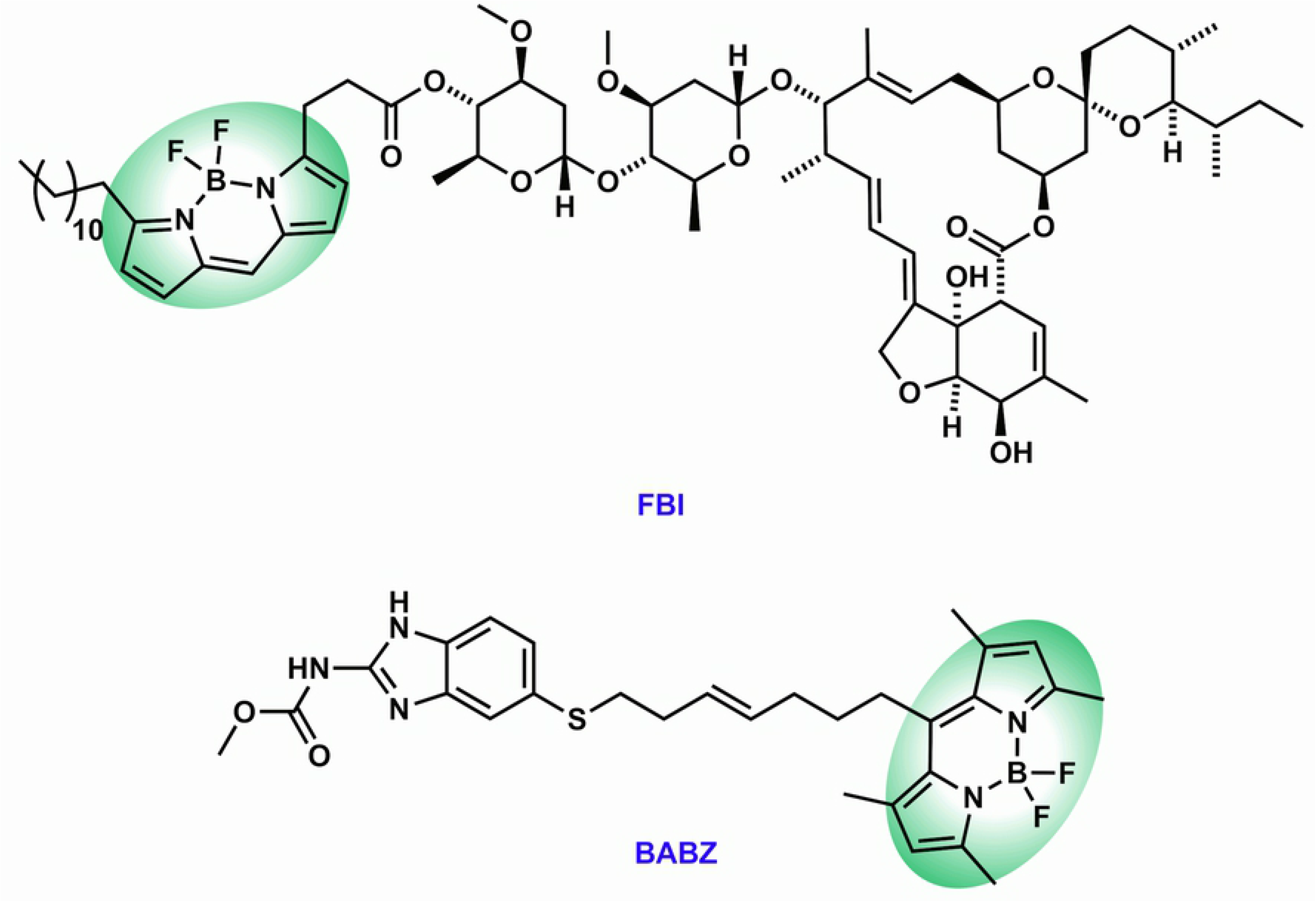
Structure of fluorescent ivermectin and albendazole analogs used. Chemical structures of the FBI and BABZ probes with the BODIPY fluorophores highlighted by a green oval.

The acute toxicity of parent anthelmintic compounds and fluorescent analogs were compared by bleaching worms to synchronise larval development before immediate use for ivermectin and FBI or rearing to L4 for albendazole and BABZ. Worms suspended in M9 were transferred to Eppendorfs in batches of 300 and made up to 184-188µl with M9 before adding 10µl OP50-1 culture and 2-6µl of anthelmintic solution (200µl total volume) and incubated for 24 hours at 21°C. The worms were then washed and split onto 3 NGM plates and for ivermectin and FBI the number of living worms were counted at 0 and every 48 hours after transfer until death or adulthood while for albendazole and BABZ all worms were immediately assessed for an uncoordinated phenotype using the same method as used to check for albendazole resistance. Ivermectin and FBI stock solutions were diluted in methanol while albendazole and BABZ stock solutions were diluted in DMSO. The range of doses tested on N2 were 10-50nM ivermectin, 1,000-15,516(1% of stock solution)nM FBI, 10-500µM albendazole and 88.45-265.35(3% stock solution)µM BABZ. For the ivermectin resistant strain DA1316, 10-3,443µM ivermectin and 9,000-15,516(1% of stock solution)nM FBI were used for the dose ranges. The albendazole resistant strain Ben-1(*e1880*) was exposed to 88.45-265.35(3% stock solution)µM BABZ. Biological replicates for each dose were performed in triplicate. Mortality and uncoordination percentages for each dose underwent a Grubb’s test for outliers (28) and was corrected against solvent only controls using the Schneider-Orelli variant of Abbott’s formula (29) before applying Probit analysis (30) to establish the lethal dose (50%)(LD_50_) for ivermectin and FBI and effective dose (50%)(ED_50_) for albendazole and BABZ. The statistical significance of LD/ED_50_ differences between strains and compounds was determined using the Litchfield & Wilcoxon method (31).

### DiI dye-filling, FBI and BABZ assays and microscopy

Worms were washed from populated plates using M9 buffer (3g KH_2_PO_4_, 6g Na_2_HPO_4_, 5g NaCl and 1mM MgSO_4_ per litre) and collected in 1.5ml Eppendorfs. Samples were pelleted by centrifugation at 7,000 rpm for 10 secs to allow removal of the supernatant. Two washes with M9 were performed before applying 10µg/ml DiI (1,1′-dioctadecyl-3,3,3′,3′-tetramethylindocarbocyanine perchlorate) dye in M9 buffer for 30 mins. Samples were then washed twice with M9 before incubating at 21°C for 2 hours to allow worms to clear their gut of bacteria and dislodge DiI adhered to the cuticle before performing two more washes in M9. Worms were pelleted and supernatant removed before transfer to an empty petri dish using a pipette and then picking 20-30 specimens onto prepared microscope slides. Slides were coated with a pad of 2% agar with 1% sodium azide and wet with 10µl of M9 containing 0.2% sodium azide, then coverslips were sealed with a thin layer of petroleum jelly.

Plates for FBI and BABZ assays were prepared by drying 3cm NGM plates in a laminar air flow cabinet for 40 mins before applying 100µl of 5µM FBI, 150µM BABZ or 150µM 1,3,5,7-tetramethyl-8-pent-4-ene-BODIPY (control for cleavage of the BODIPY containing side chain of BABZ) diluted in methanol and left for 1-3 hours prior to applying 50µl of OP50-1 culture. The next day, 40 worms of the strain to be tested were then picked onto the plates before incubation at 16°C. After the defined incubation period, 20-30 specimens were picked into droplets of M9 to wash off excess BODIPY labelled compound before picking onto prepared agarose pads on microscope slides.

Slides were viewed using a Zeiss Axioskop 2 Plus microscope fitted with a Zeiss Mercury HBO 100 Lamphouse and Zeiss AxioCam camera with images taken using the accompanying Axiovision software. Control images of worms were taken using a Differential Interference Contrast (DIC) filter, 0.5 secs exposure time and the minimum setting for the internal light source while DiI, BABZ and 1,3,5,7-tetramethyl-8-pent-4-ene-BODIPY staining was viewed and imaged using a Fluorescein Isothiocyanate (FITC) filter, 1 sec exposure time and illumination by the mercury lamp. FITC images of FBI exposed worms used a 2 sec exposure time. A minimum of 10 individuals of each strain were observed under FITC conditions to score the average intensity of DiI, FBI and BABZ staining (negative (-), weak positive (+ (w)) or strong positive (+)). Representative DIC and FITC images for DiI, FBI and BABZ staining patterns in each category are shown in Fig 2 while images for individual strains are available upon request.

**Fig 2.**
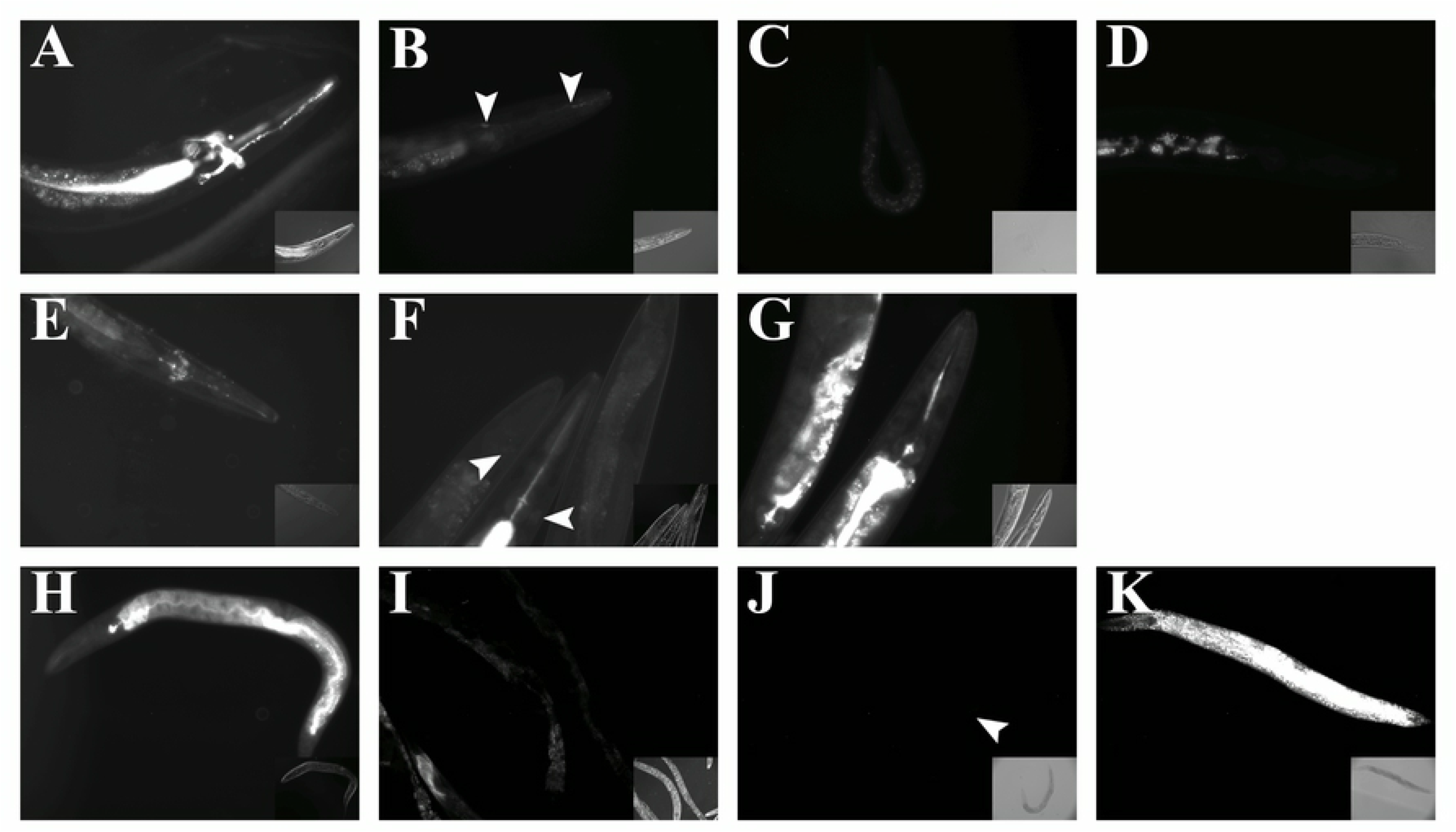
Representative images of DiI and BODIPY labelled anthelmintic analog phenotypes. DiI = 1,1′-dioctadecyl-3,3,3′,3′-tetramethylindocarbocyanine perchlorate; FBI = <u>F</u>atty-<u>B</u>ODIPY-Ivermectin; <u>B</u>ODIPY-<u>AlB</u>enda<u>Z</u>ole. (A) N2: DiI dye filling positive, (B) Ifta-1(*nx61*): Weak DiI dye filling positive, (C) Dyf-2(*m160*): DiI dye filling negative (D) C14h10.2(*tm10737*): Novel Dyf mutant which has variable DiI dye filling with weak positive individuals in a predominantly negative (pictured) population, (E) N2: FBI uptake positive, (F) Hyls-1(*tm3067*): Weak FBI uptake positive, (G) Osm-5(*p813*): FBI uptake negative, (H) N2: BABZ uptake positive, (I) Dnc-1(*or404*) Weak BABZ uptake positive, (J) Rab-35(*b1013*): BABZ uptake negative and (K) N2: Staining pattern of 1,3,5,7-tetramethyl-8-pent-4-ene-BODIPY. Individuals were photographed using a DIC filter (lower right inset image) to highlight the position and orientation of the worm and a FITC filter (main image) to visualise fluorescence. Areas of fluorescence for weak phenotypes are highlighted with arrows.

### EMS mutagenesis and whole genome sequencing

*C. elegans* L4 stage N2 strain worms were exposed to 50 mM ethyl methanesulfonate (EMS) for 4 h at 20°C following standard mutagenesis procedures (32), then allowed to recover on OP50-1 seeded NGM plates overnight. Worms were then handled according to Page, 2018 (23) selecting for 10nM moxidectin resistance (see Supplementary Methods for details). Lines were then characterised for DiI dye-filling and ivermectin, albendazole and levamisole cross resistance.

From the 14 resulting moxidectin resistant lines 5 were selected and together with uncharacterised ivermectin resistant lines TP236(ka30), TP241(ka35), TP272(ka64) and TP274(ka66) from a previous study (23) were processed for single nucleotide polymorphism (SNP) mapping. SNP mapping was carried out as described in Doitsidou, 2010 (33) using MiModD tools on the public instance of the Galaxy platform (https://usegalaxy.org)(34) (see Supplementary Methods for details). Genomic DNA was extracted using a Gentra Puregene Core Kit A (Qiagen, UK) kit before clean up and concentration using a Genomic DNA Clean & Concentrator-25 (Zymo Research, US) kit. Samples were sent for whole genome sequencing to the Glasgow Polyomics facility, University of Glasgow where libraries were prepared with a TruSeq® Nano DNA LT Sample Prep Kit (Illumina), quality controlled on a 2100 Bioanalyzer (Agilent) and run on an Illumina MiSeq platform using 300bp paired end reads.

## Results

Outcomes of dye-filling, survival and uncoordinated phenotype assays are listed in Table 1. Strains tested that did not show a phenotype of interest are included in Table S1.

**Table 1.**
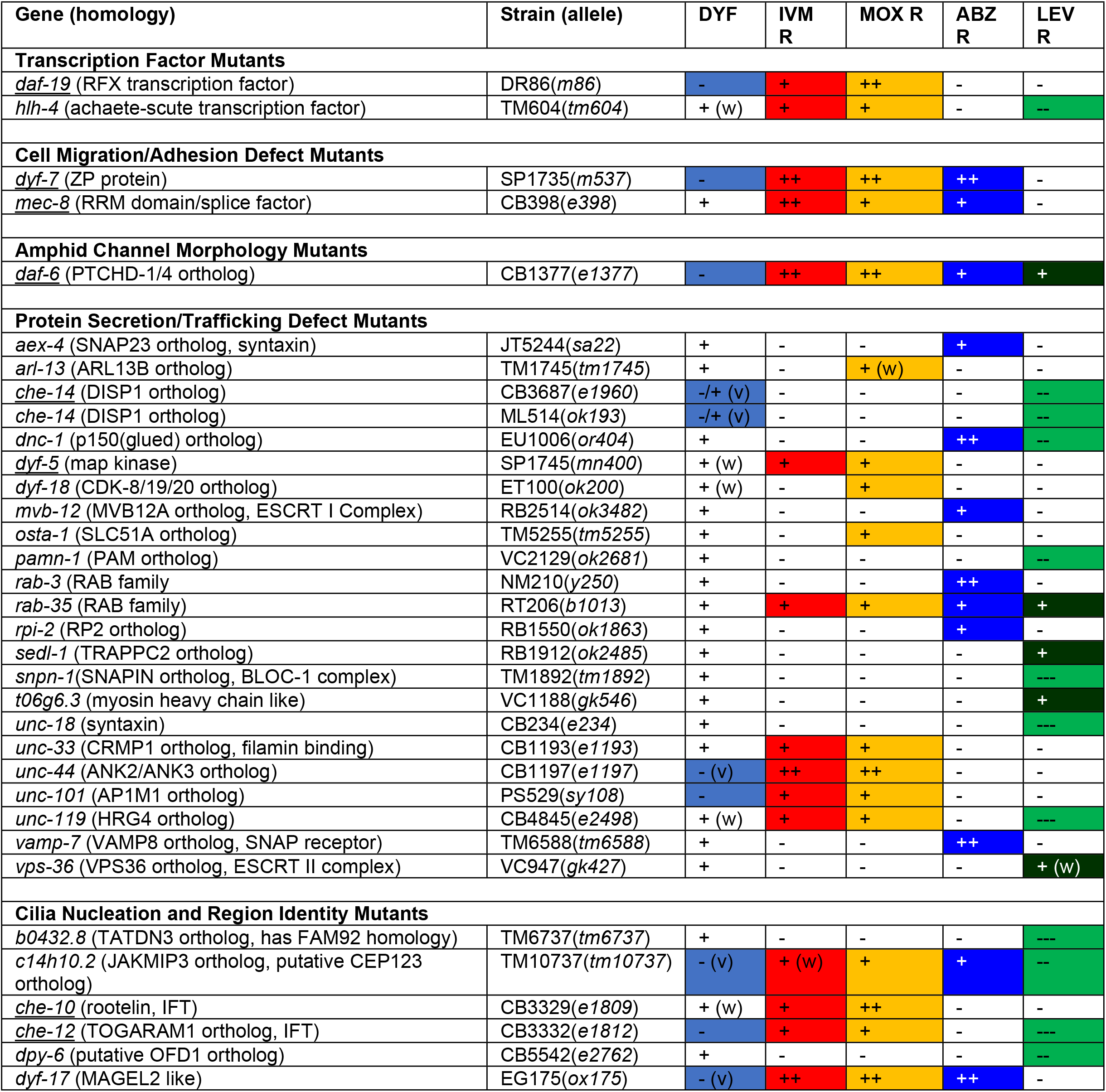
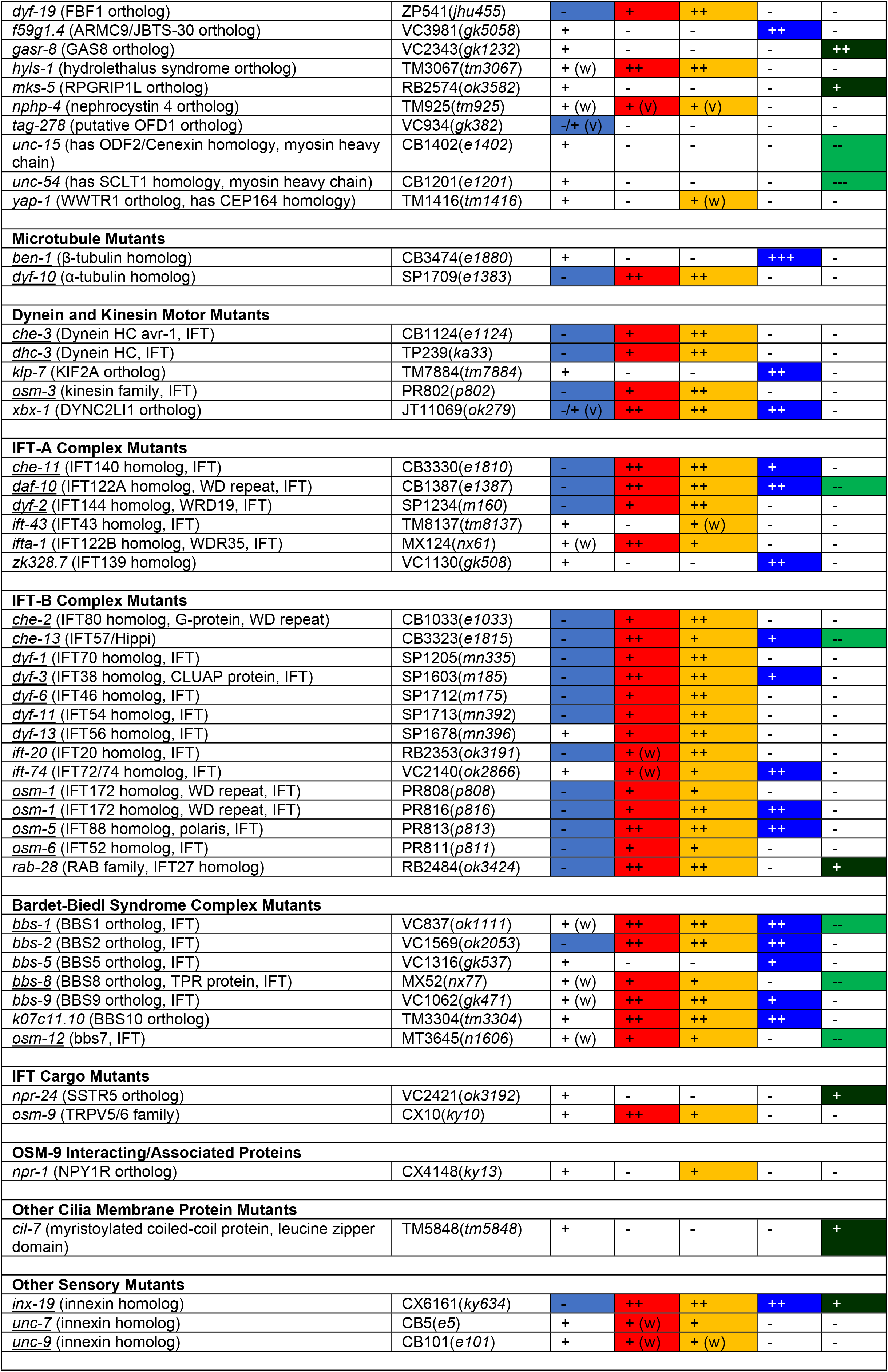
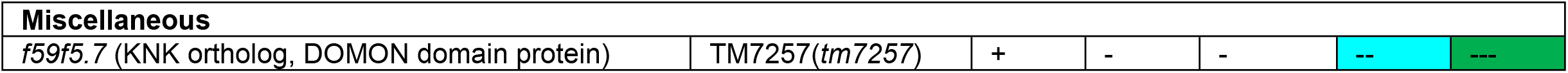
Many *C. elegans* mutants for ciliary proteins are resistant to ivermectin, moxidectin, albendazole and levamisole. **Underline** = previously identified mutants that are resistant to one of the anthelmintics tested. **Dyf** = DiI amphid dye filling; **IVM R** = Ivermectin resistance; **MOX R** = Moxidectin resistance; **ABZ R** = Albendazole resistance; **LEV R** = Levamisole resistance; **+++** = Extreme resistance; **++** = Strong resistance; **+** = Dye filling/Moderate resistance; **-** = Dye filling defective/Susceptible; **--** = Highly susceptible; **---** = Extremely susceptible; **w** = weak phenotype; **v** = High variability in phenotype (**-/+**: phenotypes range across the entire spectrum). **IFT** = intraflagellar transport component homology. *H. contortus* homologues investigated by BLASTP using WormBaseParaSite (WBPS14, WS269), hits shown as % identity over specified amino acid length.

### Intraflagellar transport complex subunits

Of the previously 34 identified ivermectin resistance genes (12, 22, 23), 16 encode for proteins of the IFT-A complex, IFT-B complex and the BBSome, all of which are interacting multiprotein complexes involved in intraflagellar transport. Therefore orthologs of the remaining 14 known, but untested, subunits of these complexes and an ortholog of the chaperone protein BBS10, were investigated for anthelmintic resistance. Out of the 15 genes tested, mutant alleles for 8 showed resistance to ivermectin. Within the IFT-A complex mutants, the IFTA-1 dynein interacting protein was found to be strongly resistant to ivermectin, while mutants for the dynein loading proteins IFT-43 and ZK328.7 remained susceptible. From the IFT-B complex mutants, the Golgi vesicle sorting protein IFT-20 and the tubulin delivery protein IFT-74 were only weakly resistant to ivermectin whereas the IFT27 ortholog RAB-28 was highly resistant. Mutants for the core BBSome proteins BBS-2 and BBS-9 and the BBS10 ortholog K07C11.10 all displayed strong ivermectin resistance while those for the cargo adapter subunits BBS-4 and BBS-5 were susceptible.

### Known IFT cargoes

As the primary function of IFT is the delivery of ciliary proteins, genes for known IFT cargo proteins were tested for ivermectin resistance to identify downstream effectors of resistance. Of the 14 cargo-protein encoding genes tested, only the CX10(*ky10*) mutant of *osm-9* was found to exhibit resistance however this finding was not replicated with the VC1262(*ok1677*) and JY190(*yz6*) *osm-9* mutant strains indicating that perhaps resistance is caused by an unrelated, uncharacterised, mutation in the CX10(*ky10*) strain. The ciliary membrane protein cargo adaptor Tub-1(*ok1972*) mutant was found to be susceptible to ivermectin, supporting the hypothesis that the downstream effector for ivermectin resistance must be delivered by another secretion pathway.

### Protein trafficking pathways

To gain insight into the trafficking of the downstream effectors for ivermectin resistance, known ciliary protein secretion pathways upstream of the IFT and ciliary membrane protein removal pathways were investigated. The clathrin adapter protein-1 ortholog involved in Golgi vesicle secretion UNC-101, the CRMP1 ortholog involved in polarizing axon-dendrite sorting UNC-33, the ANK2/ANK3 ortholog involved in polarizing axon-dendrite sorting UNC-44, UNC-119 which inserts myristoylated proteins into the cell membrane and RAB-35 which regulates early endosome recycling were all involved in ivermectin resistance. Mutants for the two SNAP25 family protein encoding genes *aex-4* and *ric-4* were found to be susceptible, supporting the contention that the downstream effector for ivermectin resistance must be delivered via vesicle fusion using the essential SNAP-29 protein. All the genes so far tested that are involved in endocytosis, designation to lysosomal degradation, early endosome maturation, extracellular vesicle formation, synaptic vesicle fusion and other post-Golgi transport complexes did not confer ivermectin resistance. Intriguingly, mutants for the RAB-8 and RAB-10 exocytosis regulators, which have roles in crossing the ciliary gate, were likewise susceptible to this drug.

Dyneins and kinesins play an important role in protein trafficking and IFT with Osm-3(*p802*), Che-3(*e1124*) and Dhc-3(*ka33*) already being associated with ivermectin resistance (12, 23), therefore additional members of these families were investigated. Of the 20 genes tested, only mutations in the dynein light-intermediate chain *xbx-1* resulted in ivermectin resistance. Mutant alleles for all three genes encoding the IFT heteromeric kinesin (*kap-1*, *klp-11* and *klp-20*) and the axonal kinesin *unc-104* had no impact on ivermectin resistance.

### The ciliary gate

The ciliary gate of the basal body acts as a physical barrier at the base of the cilia that selectively allows the passage of ciliary proteins. Components of the ciliary gate (some putative) were therefore investigated to uncover those required to deliver downstream effectors associated with ivermectin resistance. The MAGEL2 like protein DYF-17, the distal appendage interacting subunit of the basal body HYLS-1, the FBF1 ortholog DYF-19, the transition fibre subunit NPHP-4 and the JAKMIP3 ortholog with CEP123 homology C14H10.2 were all found to be involved in maintaining ivermectin susceptibility although some Nphp-4(*tm925*) individuals showed incomplete penetrance of the resistance phenotype. Mutants for all other transition fibre genes, putative subdistal appendage proteins, putative ESCRT complex, Exocyst vesicle, TRAPP complex and Rab interacting basal body subunits and orthologs of the ARMC9/TOGARAM1 complex were all tested and found to have no impact on ivermectin resistance.

### Cell migration, amphid formation, ciliogenesis and ciliated neuron enriched genes tested

As gross morphological defects to amphid neurons, their cilia and the amphid channel invariably cause ivermectin resistance, some transcription factors that determine amphid neuron cell fate and the proteins involved in axon guidance and lumen formation were assessed for a role in ivermectin resistance. Of the 5 genes tested only mutant alleles for the ADL neuron determining transcription factor *hlh-4* and the lumen endocytosis regulator *daf-6* were found to cause resistance to ivermectin.

Some genes involved in gap junction formation (*unc-7* and *unc-9*), mechanosensation (*mec-1* and *mec-8*) and osmotic avoidance (*osm-1*, *osm-3*, *osm-5*, *osm-6* and *osm-12*) cause ivermectin resistance (12, 23), so additional genes in those categories along with several cilia enriched membrane proteins (35, 36) were likewise investigated. Of the genes from this grouping that have been tested only the gap junction innexin Inx-19(*ky634*) mutant displayed resistance to ivermectin.

### Amphidal dye-filling defect correlation with ivermectin resistance

It has previously been found that there is a correlation between ivermectin resistance and dye-filling defects (23), so the full extent of this relationship was examined. Of previously known ivermectin resistance genes, the mutant alleles Daf-19(*m86*), Dyf-7(*m537*), Che-12(*e1812*), Dyf-10(*e1383*), Che-3(*e1124*), Dhc-3(*ka33*), Osm-3(*p802*), Che-11(*e1810*), Daf-10(*e1387*), Dyf-2(*m160*), Che-2(*e1033*), Che-13(*e1815*), Dyf-1(*mn335*), Dyf-3(*m185*), Dyf-6(*m175*), Dyf-7(*m537*), Dyf-10(*e1383*), Dyf-11(*mn392*), Osm-1(*p808*), Osm-1(*p816*), Osm-3(*p802*), Osm-5(*p813*) and Osm-6(*p811*) were dye-filling negative; Bbs-8(*nx77*), Che-10(*e1809*), Bbs-1(*ok1111*) and Osm-12(*n1606*) and Dyf-5(*mn400*) exhibited weak dye-filling; Che-1(*p672*), Che-1(*ot75*), Che-6(*e1126*), Dyf-13(*mn396*), Mec-1(*e1066*), Mec-8(*e398*), Unc-7(*e5*) and Unc-9(*e101*) were dye-filling positive; and Che-14(*e1960*) exhibited highly variable degrees of dye-filling between individuals. Among the novel ivermectin resistance genes identified in the present study, the mutant alleles Unc-101(*sy108*), Daf-6(*e1377*), Ift-20(*ok3191*), Rab-28(*ok3424*), Bbs-2(*ok2053*), Dyf-19(*jhu455*) and Inx-19(*ky634*) were all dye-filling negative; Hlh-4(*tm604*), Unc-119(*e2498*), Hyls-1(*tm3067*), Nphp-4(*tm925*), Ifta-1(*nx61*) and Bbs-9(*gk471*) displayed weak dye-filling; Rab-35(*b1013*), Unc-33(*e1193*), Ift-74(*ok2866*) and K07c11.10(*tm3304*) were dye-filling positive; and C14h10.2(*tm10737*), Dyf-17(*ox175*), Unc-44(*e1197*) and Xbx-1(*ok279*) had highly variable degrees of dye-filling between individuals, with C14h10.2(*tm10737*), Dyf-17(*ox175*) and Unc-44(*e1197*) being predominantly dye-filling negative. The Tag-278(*gk382*) mutant also showed highly variable degrees of dye-filling between individuals but showed no resistance to any of the tested anthelmintics. Processes that are essential for ciliogenesis and cilia maintenance showed a strong correlation between the extent of dye-filling defects and the strength of ivermectin resistance although Mec-8(*e398*), Hyls-1(*tm3067*), Ifta-1(*nx61*) and Bbs-9(*gk471*) defied the trend by showing strong resistance despite having weak dye-filling. Mutants for proteins which are involved in trafficking cilia membrane proteins along the axon such as UNC-33 and proteins which function downstream of IFT, including RAB-35 and helper/regulatory proteins like K07C11.10 showed no correlation. This indicates that although DiI dye-filling and ivermectin uptake require effector delivery to the cilia through shared pathways, both processes do not necessarily use the same effector.

### Observed resistances to other anthelmintics and roles of IFT in cross resistance

Candidate genes were also tested for moxidectin (an avermectin), albendazole and levamisole (both non-avermectins) resistance to examine possible cross resistance. Mutants for all genes that were ivermectin resistant were also resistant to moxidectin, indicating as expected, shared mechanisms but also similar levels of resistance to the two drugs. Mutants for the kinase DYF-18 which plays a role in ciliogenesis and IFT, a regulator of ciliary protein trafficking OSTA-1, the small GTPase nucleotide exchange factor involved in ciliogenesis ARL-13, a WWTR1 ortholog with CEP164 homology YAP-1 and the IFT-A complex dynein loading protein IFT-43 however, showed moxidectin resistance but not ivermectin resistance. The CX4148(*ky13*) mutant strain of the OSM-9 interacting protein NPR-1 showed moxidectin resistance, however this finding was not reproduced for a second allele using the RB1330(*ok1447*) strain indicating that it is most probably caused by an unrelated, uncharacterised, mutation in the CX4148(*ky13*) strain.

Cross resistance to the unrelated benzimidazole drug albendazole was observed for genes involved in cell migration and IFT with *daf-6*, *dyf-17*, *rab-35*, *c14h10.2* and *inx-19* also showing cross resistance. Within the IFT-B complex, resistance was limited to the tubulin interacting protein IFT-74, the BBSome interacting DYF-3/CHE-13 dimer and the SEC-24 (COPII ortholog) pathway interacting protein OSM-5. The BBSome cargo adapter proteins which are important for albendazole susceptibility differed from those for avermectins with Bbs-5(*gk537*) mutants showing resistance while Bbs-8(*nx77*) mutants were susceptible. Several other genes whose mutation confers albendazole resistance were identified including the IFT-A complex dynein loading protein *zk328.7*, the SNAP25 family member *aex-4*, the VAMP8 ortholog SNAP receptor *arl-7*, the SEC-24 (COPII ortholog) pathway interacting p150^glued^ ortholog *dnc-1*, the ARL-3 activating kinase *rpi-2*, the negative regulator of microtubule length *f59g1.4*, the ESCRTI complex subunit *mvb-12*, the regulator of vesicle trafficking for endocytosis and exocytosis *rab-3*, and the kinesin *klp-7*. Of all genes tested only the KNK ortholog *f59f5.7* was found to have increased albendazole susceptibility showing a greatly reduced population growth, in comparison to controls, that was unable to clear the plates of OP50-1 within 144 hours. None of the mutants tested showed resistance as strong as the well characterised albendazole resistant mutant control Ben-1(*e1880*) (37).

Levamisole, represents the third unrelated class of anthelmintic examined for cross resistance. Levamisole resistance and susceptibility had no obvious connection to the other anthelmintics tested, however Rab-35(*b1013*), Daf-6(*e1377*) and Inx-19(*ky634*) mutants displayed moderate broad-spectrum cross resistance and Rab-28(*ok3424*) had resistance to both avermectins and levamisole. The uncharacterised gene T06g6.3(*gk546*), the TRAPP-I/II complex subunit Sedl-1(*ok2485*), nexin link protein Gasr-8(*gk1232*), transition fibre subunit Mks-5(*ok3582*), the IFT cargo Npr-24(*ok3192*), the amphid exosome export protein Cil-7(*tm5848*) and the ESCRT-II complex subunit Vps-36(*gk427*) mutants were however all found to confer resistance to levamisole. Mutants for Bbs-1(*ok1111*), Bbs-8(*nx77*), Che-13(*e1815*), Che-14(*e1960*), Dnc-1(*or404*), Daf-10(*e1387*), the putative OFD1 ortholog Dpy-6(*e2764*), Hlh-4(*tm604*), Osm-12(*n1606*), the PAM ortholog Pamn-1(*ok2681*), C14h10.2(*tm10737*), and the myosin heavy chain with cenexin homology Unc-15(*e1402*) all showed greatly reduced population growth compared to controls and the other tested susceptible strains, indicating an increased susceptibility to levamisole. Although levamisole exposure usually does not kill *C. elegans*, even at doses as high as 10mM (determined from preliminary dose ranging experiments using N2), high mortality was observed in the syntaxin Unc-18(*e234*), the BLOC1 complex subunit Snpn-1(*tm1892*), the uncharacterised protein with FAM92 homology B0432.8(*tm6737*), the myosin heavy chain with SCLT1 homology Unc-54(*e1201*), Unc-119(*e2498*), the TOGARAM1 ortholog Che-12(*e1812*) and F59f5.7(*tm7257*) mutants at the 0.2mM and 0.8mM doses used for resistance assays.

### Visualisation of anthelmintic uptake using BODIPY labelled analogs

To visualise the route of ivermectin and albendazole uptake, chemical analogs which were linked to a BODIPY fluorophore were constructed and applied to live worms. The ivermectin probe was named FBI and the albendazole probe called BABZ. It was confirmed that the analogs retained some of their target binding using acute toxicity assays to compare the LD/ED_50_s of analogs to the parent compounds in sensitive N2 and target site insensitive DA1316, for ivermectin and FBI, and Ben-1(*e1880*), for albendazole and BABZ, backgrounds. Ivermectin was found to have an LD_50_ of 25.98nM (95%CI = 24.49-27.56 nM; N = 3,330) for N2 and 496.33µM (95%CI = 432.52-569.56µM; N = 8,024) for DA1316 while FBI had an LD_50_ of 6.35 µM (95%CI = 5.76-7 µM)(N = 2,098) for N2 and 83.59µM (95%CI = 35.43-197.18µM; N = 1,996) for DA1316. This indicates the primary toxicity, caused by target binding, of FBI is 244 times lower than the parent compound while secondary toxicity, caused by off target effects, is 5.9 times higher. The ED_50_ for albendazole was found to be 51.63µM (95%CI = 43.71-60.98µM; N = 3,607) for N2 and estimated to be 1.68mM (95%CI = 1.53-1.85mM; N = 2,492) for Ben-1(*e1880*) while the ED_50_ for BABZ was estimated to be 252.28µM (95%CI = 216.96-293.35µM; N = 3,307) for N2 and 471.39µM (95%CI = 424.18-523.85µM; N = 1,646) for Ben-1(*e1880*). This suggests that the primary toxicity of BABZ is 4.9 times lower than the parent compound while secondary toxicity is 3.6 times higher.

Time course experiments were performed on N2 worms to characterise uptake progression and establish the optimum incubation time with the probes (see Supplementary Results and Supplementary Figs S1 and S2 for details). FBI uptake was found to be restricted to the amphid neurons with limited spread to the adjacent nerve ring while BABZ was primarily absorbed in the hind gut but showed progressive systemic spread. A 72 hour incubation period was selected to be used for all subsequent assays.

The uptake of the ivermectin probe FBI and the albendazole probe BABZ in different genotypes was then examined (Table 2). The susceptible strain CB4856 and target site insensitivity mutants DA1316 and Ben-1(*e1880*) stained in an identical manner to N2 indicating no differences in uptake between those strains. Next a selection of resistant mutants from the moxidectin forward genetic screen and the targeted resistance screen were exposed to FBI and/or BABZ, depending on the resistances of the strain, to determine if resistance is being caused by changes in flux. Uptake of FBI by the amphids occurred in Rab-35(*b1013*), Unc-7(*e5*) and Unc-9(*e101*) but not in Daf-6(*e1377*), Dyf-19(*jhu455*), Inx-19(*ky634*), Osm-5(*p813*) Osm-9(*ky10q*), TP236(*ka30*), TP241(*ka35*), TP272(*ka64*), TP274(*ka66*) and TP388(*ka204*) worms while Hyls-1(*tm3067*), Unc-44(*e1197*) and TP378(*ka201*) showed weak uptake. The distribution of absorbed BABZ in most tested resistant strains was limited to the gut with Dnc-1(*or404*), Osm-5(*p813*), TP272 and TP384 showing weak uptake while Daf-6(*e1377*), Inx-19(*ky634*), Rab-35(*b1013*) and TP386 showed almost no uptake. TP236 and TP375 were exceptions in that weak BABZ uptake was observed while still retaining systemic spread throughout the whole worm. TP241 showed positive uptake comparable to the controls suggesting the observed resistance to albendazole may be caused by target site insensitivity or enhanced phase I detoxification.

**Table 2.** Uptake patterns of BODIPY labelled anthelmintic analogs across different strains. **FBI** = Fatty-BODIPY-Ivermectin; **BABZ** = BODIPY-Albendazole; **+** = Positive uptake; **+(w)** = Weak uptake; **-** = No or barely visible uptake; **NT** = Not tested.

| Strain | N2 | CB4856 | DA1316 | Ben-1 | Rab-35 | Daf-6 | Inx-19 | Dyf-19 | Osm-5 | Dnc-1 | Osm-9 | Hyls-1 |
| --- | --- | --- | --- | --- | --- | --- | --- | --- | --- | --- | --- | --- |
| FBI | + | + | + | + | + | - | - | - | - | NT | - | +(w) |
| BABZ | + | + | + | + | - | - | - | NT | +(w) | +(w) | NT | NT |
| Strain | Unc-7 | Unc-9 | Unc-44 | TP236 | TP241 | TP272 | TP274 | TP375 | TP378 | TP384 | TP386 | TP388 |
| FBI | + | + | +(w) | - | - | - | - | - | +(w) | - | - | - |
| BABZ | NT | NT | NT | +(w) | + | +(w) | + | +(w) | + | +(w) | - | + |

### Whole genome sequencing of mutants from forward genetic screens

Extensive EMS genetic screens for ivermectin and abamectin resistant mutants were carried out previously and limited mapping identified two IFT-related mutants (Che-3(*ka32*) and Dhc-3(*ka33*)) (23). In this current study, a new forward genetic screen to identify moxidectin resistant strains was performed. Together these screens identified 31 mutants resistant to avermectins which also had their albendazole and levamisole resistance and DiI dye-filling phenotypes characterised (Table 3). Based on phenotype TP236(*ka30*), TP241(*ka35*), TP272(*ka64*) and TP274(*ka66*) from the previous abamectin screen along with TP375(*ka200*), TP378(*ka201*), TP384(*ka202*), TP386(*ka203*), and TP388(*ka204*) from the current moxidectin screen were selected for backcrossing, whole genome sequencing and SNP mapping. Of the selected strains all were resistant to ivermectin and moxidectin with TP272(*ka64*), TP274(ka66) and TP384(*ka202*) having cross resistance to levamisole and TP241(*ka35*), TP375(*ka200*) and TP386(*ka203*) having cross resistance to albendazole and levamisole with TP375(*ka200*) showing strong cross resistance to all three. TP388(*ka204*) was dye-filling positive while all others were dye-filling negative. The whole genome sequencing and mapping data (aligned reads available at https://www.ncbi.nlm.nih.gov/sra/PRJNA768320) identified novel alleles of *osm-3*, *che-3* (4 different alleles), *osm-1*, *dhc-3*, *dyf-2* and *ifta-1* (Fig 3) as the causative genes for resistance to avermectins. Details of identified alleles are listed in Table S2.

**Fig 3.**
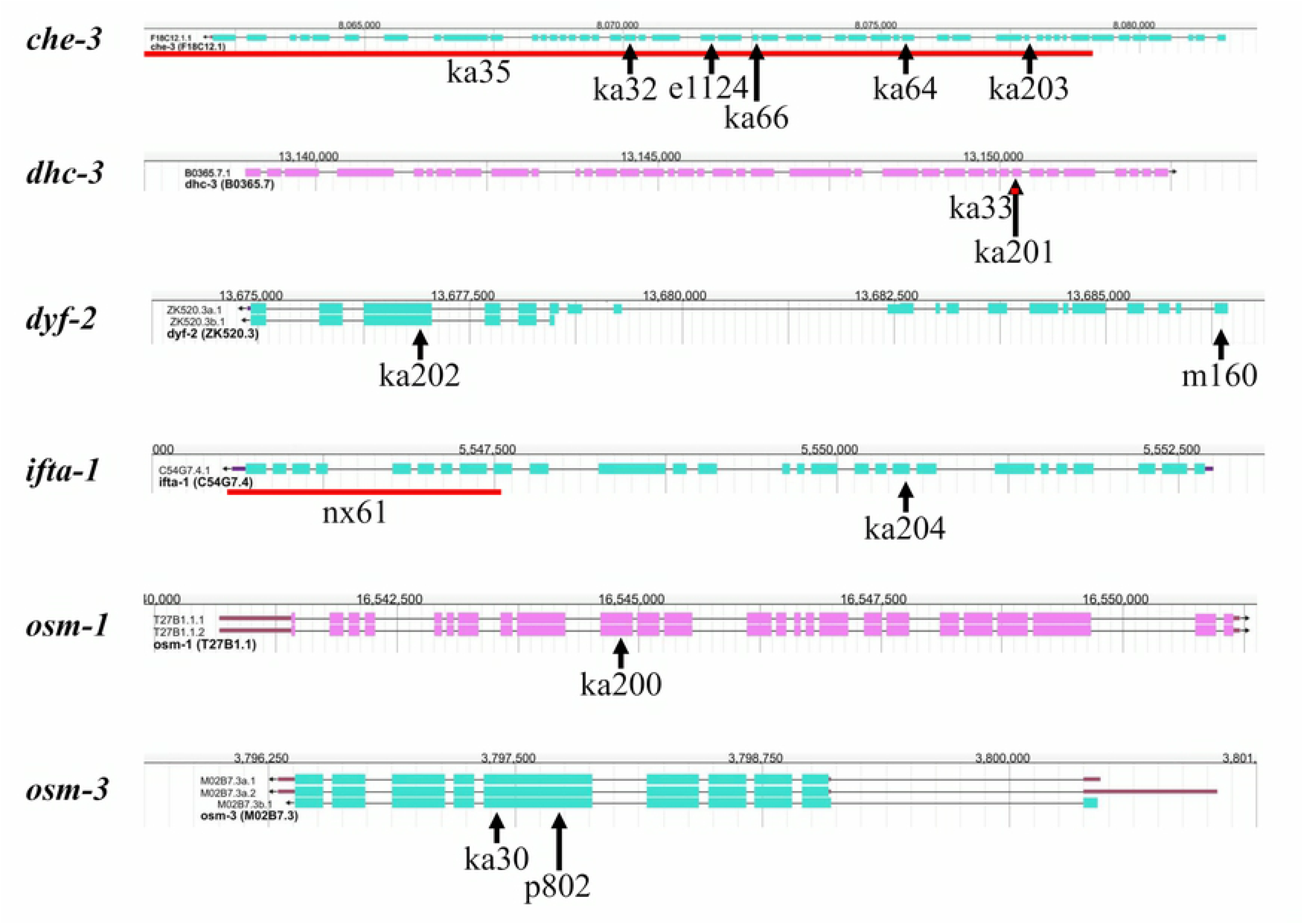
Position of novel and tested alleles in resistance genes identified by whole genome sequencing. Transcript structures and positions of genes were obtained from WormBase (https://wormbase.org) (JBrowse version: WS281; genome build WBcel235). Arrows above alleles point to their location in the genomic sequence. Red lines above alleles span the length of deletions. Alleles featured (*name* = chr-number: position nt-change (aa-change)) are ***e1124*** = I: 8,071,718 G>A (Q>Stop); ***ka30* =** IV: 3,797,404 G>A (Q>Stop); ***ka32*** = I: 8,070,133 C>T (G>R); ***ka33*** = V: 13,150,172-13,150,276 deletion; ***ka35*** = I: 8,058,869-8,079,083 deletion; ***ka64*** = I: 8,075,488 A>T (L>Stop); ***ka66*** = I: 8,072,572 C>T (E>K); ***ka200*** = X: 16,544,813 C>T (Q>Stop); ***ka201*** = V: 13,150,224 AGG>AG frameshift; ***ka202*** = III: 13,676,892 G>A (Q>Stop); ***ka203*** = I: 8,077,873 G>A splice site acceptor change; ***ka204*** = X: 5,550,502 A>T (C>Stop); ***m160*** III: 13,686,367 G>A (R>Stop); ***nx61* =** X: 5,545,532-5,547,540 deletion; ***p802*** = IV: 3,797,722 G>A (Q>Stop); ***p808*** = X: Uncharacterised; ***p816*** = X: Uncharacterised ∼600bp deletion.

**Table 3.** Resistance profiles and causative genes for resistance to avermectins in EMS generated mutant strains. **Dyf** = DiI amphid dye filling; **IVM R** = Ivermectin resistance; **MOX R** = Moxidectin resistance; **ABZ R** = Albendazole resistance; **LEV R** = Levamisole resistance; **++** = Strong resistance; **+** = Dye filling/Moderate resistance; **-** = Dye filling defective/Susceptible.

| Strain (selection screen used for isolation) | Assigned allele | Causal gene | Mutation/effect | DYF | IVM R | MOX R | ABZ R | LEV R |
| --- | --- | --- | --- | --- | --- | --- | --- | --- |
| TP236 (10nM Ivermectin) | *ka30* | *osm-3* | Substitution/Nonsense | - | ++ | + | - | - |
| TP241 (50nM Abamectin) | *ka35* | *che-3* | Deletion/Coding | - | ++ | + | + | + |
| TP272 (10nM Ivermectin) | *ka64* | *che-3* | Substitution/Nonsense | - | ++ | + | - | + |
| TP274 (10nM Ivermectin) | *ka66* | *che-3* | Substitution/Missense | - | ++ | + | - | + |
| TP375 (10nM Moxidectin) | *ka200* | *osm-1* | Substitution/Nonsense | - | + | ++ | ++ | ++ |
| TP378 (10nM Moxidectin) | *ka201* | *dhc-3* | Deletion/Frameshift | - | ++ | ++ | - | - |
| TP384 (10nM Moxidectin) | *ka202* | *dyf-2* | Substitution/Nonsense | - | + | ++ | - | + |
| TP386 (10nM Moxidectin) | *ka203* | *che-3* | Splice site substitution | - | ++ | ++ | + | + |
| TP388 (10nM Moxidectin) | *ka204* | *ifta-1* | Substitution/Nonsense | + | ++ | ++ | - | - |

## Discussion

### IFT protein resistances and redundancies

IFT is highly conserved throughout eukaryota, being required for the import and transport of ciliary proteins to their correct localisations within the cilia (38) (Fig 4A). Loss of IFT impacts on cell motility, cell migration, cell signalling, cell division and the ability to sense environmental stimuli with mutants for mammalian orthologs of IFT proteins being responsible for 16 of the 35 known disease causing ciliopathies (38–40). IFT mutants have also been recently linked to ivermectin resistance in *C. elegans* (12, 22, 23). The protein-protein interactions of the IFT-A and IFT-B complexes (38, 41–46) and the BBSome (46–48) are well documented however not all interactions have been verified in *C. elegans*. When the identified resistance causing genes are overlaid with known and predicted interactions (Fig 4B) potential mechanisms for resistance become apparent.

**Fig 4.**
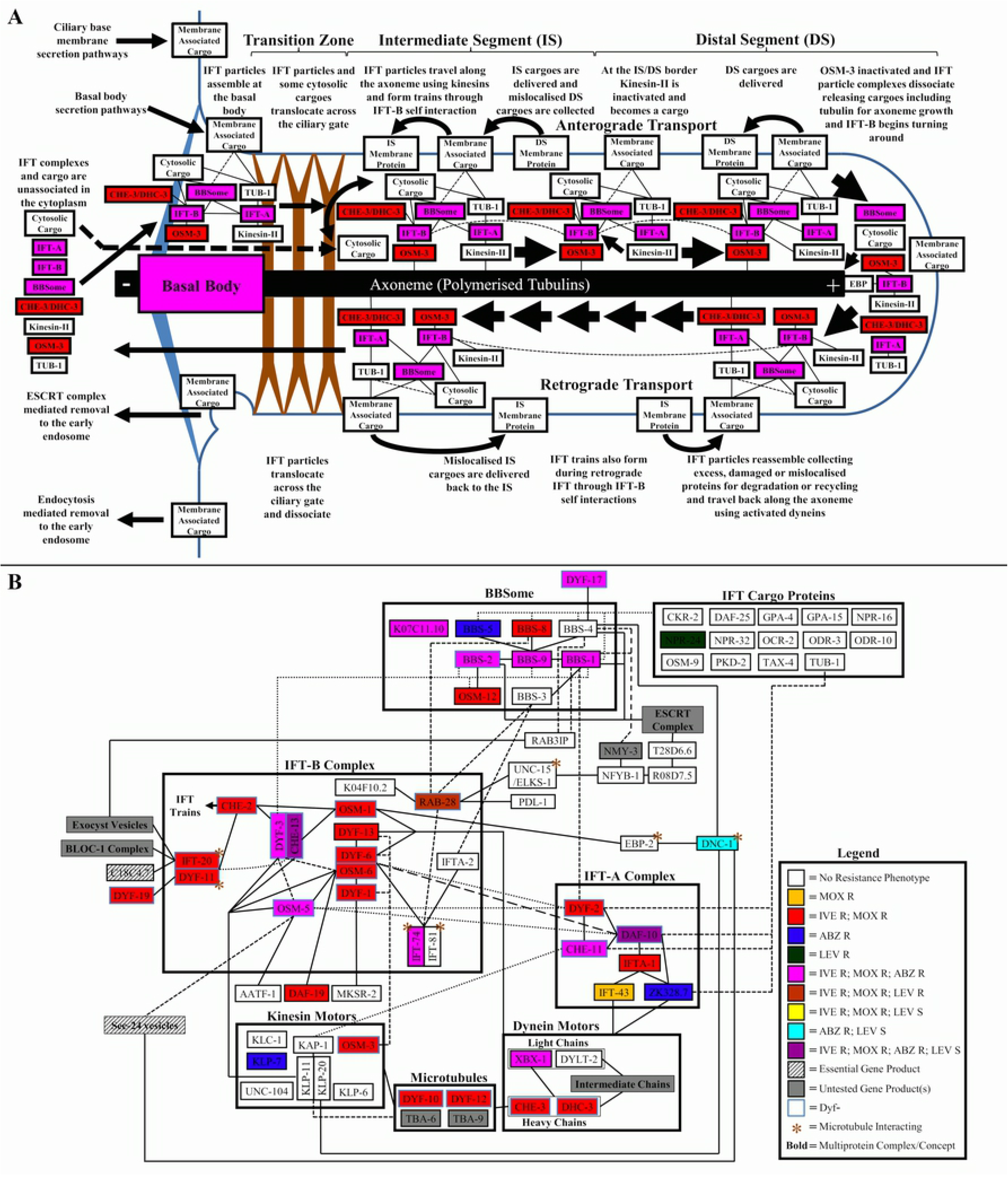
Intraflagellar transport in *C. elegans* and resistance patterns in the IFT protein-protein interaction network. (A) Summary of *C. elegans* intraflagellar transport. Colours used are for the summation of resistances found in complexes. **IFT-A** = Intraflagellar transport complex A; **IFT-B** = Intraflagellar transport complex B; **BBSome** = Bardet-Biedl Syndrome complex; **Line** = protein/complex-protein/complex interaction; **Small Arrow** = Change in protein or complex localisation or interaction; **Large Arrow** = Direction of IFT particle travel. (B) Predicted IFT protein-protein interaction network in *C. elegans* showing resistances found in mutants of each node. **Box** = Group of proteins from the same complex or with the same function; **Line** = predicted protein-protein interaction; **Small Arrow** = Protein self-interaction; **/** = Multiple candidate genes with homology to a node found in other species.

In *C. elegans* the kinesins responsible for anterograde IFT are redundant in the intermediate segment of the cilia, with both the homomeric kinesin OSM-3 and the heteromeric kinesin (KAP-1, KLP-11 and KLP-20) being sufficient to build and maintain the intermediate segment. The distal segment however, is dependent solely on OSM-3 function (38). Mutants for the heteromeric kinesin had no impact on ivermectin resistance while those for *osm-3* were resistant (23) indicating that the downstream effectors for ivermectin susceptibility are transported to the distal segment of the cilia. There is further indication that the effector protein localises to the distal segment through analysis of the Unc-101(*sy108*) and Unc-119(*e2498*) mutants, both of which lack distal segments (49) and the Dyf-5(*mn400*) and Dyf-18(*ok200*) mutants which have elongated middle segments that invade the distal segment (50–52), since all these mutants show resistance. There is evidence that the downstream effector for ivermectin resistance needs to be transported back to the base of the cilia by retrograde IFT for ivermectin susceptibility, since mutation of the ciliary dynein heavy chains CHE-3 and its paralogue DHC-3 together with their interacting light-intermediate chain XBX-1 (53) cause resistance. The importance of effector protein retrieval from the cilia for ivermectin susceptibility is highlighted by the resistance seen in mutants for DYF-6, OSM-1 and DYF-13 which are involved in dynein import into cilia and the turning around of IFT complexes at the distal tip (54–57).

The cilia axoneme is composed of polarised tubulins (including DYF-10 and DYF-12 (58)) that grow distally from the basal body and require a high local concentration of tubulins to polymerise, confirming the requirement for their delivery to the tip of the cilium (59). Tubulin heterodimers are imported into the cilia via the redundant IFT-B proteins IFT-74 and IFT-81 (60). Consequently Ift-74(*ok2866*) mutants only show weak resistance to ivermectin indicating that unlike other species, the *C. elegans* dimer subunits, do not have an equal role in tubulin import. The distance that the IFT cargoes are transported on the axoneme is determined by the stability of the interaction between the complexes and the microtubule rail, with the ability to form IFT trains through IFT-80 (CHE-2) interaction (44) and the microtubule interacting DYF-11/IFT-20 dimers playing an important role in this process (61). The interactions of the three gene products is reflected in their shared ivermectin resistance, while the low level of resistance observed in IFT-20 mutants can be attributed to a known inequality in functional redundancy of the two heterodimer subunits (61). The tubulin interacting IFT-B subunit IFT-74 was also found to play a role in albendazole resistance, potentially through the transport of poisoned BEN-1 subunits to the plus end of microtubules or delivery of unexposed BEN-1 to the interface with albendazole. The OSM-6/DYF-6 dimer physically bridges between the two IFT-B core subcomplexes that contain the albendazole resistance causing subunits IFT-74 and OSM-5, DYF-3 and CHE-13 respectively (62). The OSM-6/DYF-6 dimer is also essential for IFT-B targeting to the basal body and for stability during transport (63–65) and mutants are susceptible to albendazole indicating either the dimer subunits have a redundancy in *C. elegans* or the IFT-B interactions with albendazole are occurring outside of IFT.

RAB-28 is a prenylated protein that functions both as an IFT27 ortholog and a GTPase, with mutants causing dye-filling defects through amphid pore malformations (66, 67). As the interaction of RAB-28 with IFT is dependent on PDL-1 mediated prenylation and BBS-3 interaction (66, 68), the results suggest that the ivermectin resistance observed in the *rab-28* mutant is being caused by the BBS-8 mediated interaction with the periciliary membrane (66, 67). The farnesylated-protein converting enzymes FCE-1 and FCE-2 process proteins for prenylation (69). FCE-1 and FCE-2 play no role in ivermectin resistance and therefore it is likely that the downstream effectors and proteins for all essential ciliary protein delivery pathways do not require prenylation to function. In *Chlamydomonas reinhardtii* BBS-3 is known to interact with another IFT-B subunit, IFT-22 (IFTA-2) (70), where they both play an important role for BBSome recruitment, however evidence from our results suggest that this interaction either does not occur in *C. elegans* or is not essential for IFT as neither gene had a role in dye-filling or ivermectin resistance.

Within the BBSome, loss of function induced ivermectin resistance was observed in DYF-3 in the IFT-B complex which interacts with OSM-12 (71) and across the core BBSome proteins (BBS-1, BBS-2, BBS-9 and OSM-12) to the BBS-1 interacting protein in the IFT-A complex, DYF-2 (72). This indicates the important role the BBSome plays in bridging the IFT-A and IFT-B complexes during IFT. The BBSome acts as a carrier for protein cargoes by both delivering proteins to the correct location in the cilia and by retrieving ciliary membrane proteins for recycling or degradation (73). During transport, the IFT cargoes interact with subsets of BBS-1, BBS-4, BBS-5 and BBS-8 (74). As BBS-8 is important for ivermectin susceptibility (23) it can be deduced that the downstream effector for ivermectin susceptibility is probably an IFT transported BBSome cargo which interacts with one or more of the adapter subunits, however the role of BBS-4 and BBS-5 was not clearly defined, a fact that may relate to their functional redundancy for some cargoes (75). In the case of albendazole susceptibility, a strong interaction between the BBSome and IFT-A and IFT-B complexes does not seem to be required as Dyf-2(*m160*) and Osm-12(*n1606*) mutants are both susceptible to this drug, perhaps indicating redundancy in their interactions and the BBSome having a role upstream of IFT.

Core subunits within the IFT-A complex showed ivermectin resistance up to the dynein motor interacting IFTA-1 subunit. Mutants for the IFTA-1 interacting dynein docking proteins ZK328.7 and IFT-43 were susceptible, an observation explained by the known redundancy these proteins have when interacting with specific dyneins (45, 76, 77). Moxidectin resistance shows a similar pattern to ivermectin resistance except the resistance seen with IFT-43 loss which indicates an inequality in functional redundancy with ZK328.7. Interestingly, albendazole susceptibility did not require IFTA-1, however ZK328.7 was necessary for susceptibility indicating that ZK328.7 interacts with other core IFT-A subunits in *C. elegans*. Such interactions have been supported by other studies (77) and ZK328.7 is probably interacting via the IFT-144 ortholog DYF-2 (45).

Benzimidazoles, such as albendazole, function by binding to the colchicine-binding domain of β-tubulins resulting in the premature capping of microtubules leading to microtubule depolymerisation and a loss of cellular structure (78). BEN-1 is the only albendazole sensitive tubulin in *C. elegans* and acts redundantly with other β-tubulins in microtubule formation (37). This redundancy explains why loss of function alleles like Ben-1(*e1880*) are resistant to albendazole without causing morphological defects or cross resistances to the other anthelmintics tested. There is also evidence that BEN-1 is not a true ciliary tubulin (79) suggesting that the albendazole resistance observed in the IFT and BBSome complex mutants are occurring through a loss of interaction with BEN-1 outside of the cilia and potentially, outside the ciliated neurons.

### Secretion pathways used by cilia proteins and the ciliary gate

Cilia proteins are delivered via the SEC-24(COPII), TRAPPII, ESCRT, exocyst, BLOC-1 and PDL-1(PDE6D)/UNC-119 secretion pathways (80–83) (summarised in Fig 5A) with many being secreted from the endoplasmic reticulum via Golgi vesicles in a clathrin adapter protein-1 (UNC-101) dependent pathway (84). These proteins would include those for ciliogenesis, IFT and the downstream effectors for ivermectin susceptibility thereby providing an explanation for the ivermectin resistance seen in Unc-101(*sy108*) mutants. The SEC-24(COPII) pathway is an essential pathway making it difficult to probe directly, however orthologs of Osm-5(*p813*) and Dnc-1(*or404*), which are both resistant and showed reduced labelled albendazole (BABZ) uptake, are known to interact with the vesicle coat proteins of this pathway (80, 85) suggesting a role in albendazole susceptibility. As Osm-5(*p813*) and Dnc-1(*or404*) showed reduced but not abolished BABZ fluorescence compared to the wild type there is an indication for either redundancy in the functions of OSM-5 and DNC-1 or the existence of additional routes for albendazole uptake. In multicellular organisms the ESCRT complexes are essential for development due to their role in controlling cell surface receptor populations through facilitating endocytosis, endosome maturation and fusion of vesicles to the lysosome and by having direct roles in establishing cell polarity and cleavage during cell division (86, 87) meaning that only non-essential subunits could be investigated for roles in anthelmintic resistance. The ESCRT-I complex is known to have a genetic interaction with the BBSome (88) and both are involved in the removal of ubiquitylated receptors (86, 89) so the albendazole resistance seen in the ESCRT-I complex subunit Mvb-12(ok3482) indicates that the ESCRT pathway may be functioning downstream of the BBSome to facilitate albendazole uptake and the effector may be a monoubiquitinated protein. The results associated both the ESCRT-II complex subunit VPS-36 and TRAPP complex subunit SEDL-1 with levamisole resistance indicating that this compound may be gaining entry through multiple routes. Mutation in the exocyst pathway genes *exoc-7* and *exoc-8* have been shown to cause weak levamisole resistance during acute exposure (90) so it was surprising that these phenotypes were unable to be replicated in mutants for any of the exocyst complex genes tested (*exoc-7*, *exoc-8* and *sec-6*) indicating this pathway only plays a minor role during chronic exposure. Despite having a known interaction with IFT-20 (82), the BLOC-1 complex was not associated with survival against any of the anthelmintics tested. The UNC-119 secretion pathway of myristoylated and laurylated acyl-anchored membrane proteins (91) is known to deliver ARL-3 and ARL-13 to the cilia facilitating regulation of the assembly and disassembly of the IFT complexes (92). The observed resistance to avermectins in the Unc-119(*e2498*) and Arl-13(*tm1745*) mutants may therefore be explained by the impaired delivery of proteins involved in cilia maintenance resulting in truncated cilia (49).

**Fig 5.**
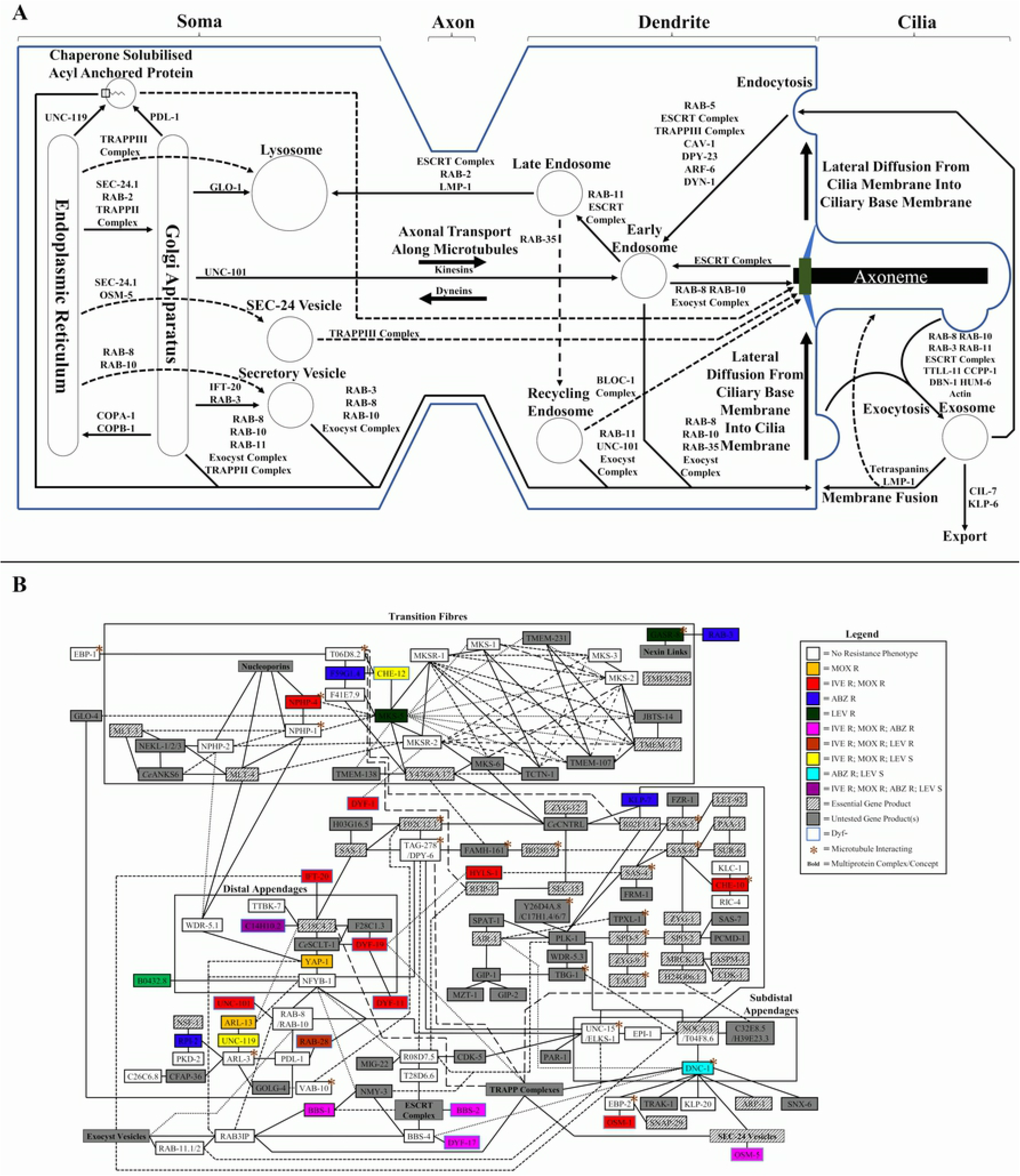
Ciliary protein trafficking pathways in C. elegans and resistance patterns in the ciliary gate protein-protein interaction network. (A) Protein trafficking pathways used to deliver and remove ciliary proteins. **Small Arrow** = Show directionality of protein trafficking between cellular locations or organelles with key proteins and complexes involved in trafficking listed next to the arrow (placed before junctions if merging into a common secretion pathway); **Large Arrow** = Directionality of axonal transport or passive diffusion. (B) Predicted basal body protein-protein interaction network in *C. elegans* showing resistances found in mutants of each node. **Box** = Group of proteins from the same complex or with the same function; **Line** = predicted protein-protein interaction; **/** = Multiple (2–4) candidate genes with homology to a node found in other species (if gene IDs differ only by the last digit, then only the last digit is shown to the right of the candidate with a similar ID); ***Ce*(node name of vertebrate ortholog)** = Multiple (>4) candidate genes with homology to the node found in other species.

The ciliary gate of the basal body (Fig 5B) acts as an impermeable barrier to proteins and macromolecules over 70 kDa in size (93) meaning that cargo delivery pathways for ciliary proteins need to interact with the basal body. To allow passage of the IFT-B complex, along with associated proteins and complexes, through the basal body MKSR-2 interacts with DYF-1 (94), DYF-19 with DYF-11 (95) and the CCDC41 ortholog C18C4.7 with IFT-20 (96). The BBSome gains entry to the cilia by interacting with the distal appendage interacting protein NMY-3 (Dzip1 ortholog)(97) while the subdistal appendage component T04F8.6 (ninein ortholog) allows passage of the dynactin complex associated cargo via the interaction with the p150^glued^ ortholog DNC-1 (98) and through cargo delivered by the exocyst pathway (99). Vesicles delivered by the exocyst pathway originate from the UNC-101 secretion pathway and have their entry regulated by RAB-8 and RAB-10 GTPases (100, 101). The RAB-8 and RAB-10 GTPases in turn localize to the basal body via interaction with cenexin and the CBY1 ortholog NFYB-1 (102, 103). The ESCRT complex mediated cargoes are potentially delivered via T28D6.6, as orthologs (DRG1 and SPI1) interact with both centrin from the basal body and the ESCRT-III complex (104, 105). Of all these gating proteins only those from the distal appendage (DYF-19 and possibly the essential protein C18C4.7) are important for resistance to avermectins indicating that the downstream effector for resistance to avermectins is delivered into the cilia as part of IFT particles and not independently via the exocyst or ESCRT secretion pathways. The protein DYF-19 is known to play an important role in facilitating the passage of IFT components across the transition zone of the basal body (95) a fact that explains why both Dyf-19(*jhu455*) mutants and those for Hyls-1(*tm3067*), which connects DYF-19 to the mother centriole of the basal body (106, 107), were also found to be resistant to avermectins and showed impaired uptake of FBI. YAP-1 is an ortholog of a transcription factor from the Hippo pathway with a role in cell cycle regulation, thermotolerance and neuronal development (108, 109) however it also has homology with the distal appendage protein CEP164 meaning that the cause of the weak moxidectin resistance observed in Yap-1(*tm1416*) needs further investigation as it could be the result of either changes in amphid neuron or cilia morphology or the upregulation of stress response pathways. The predicted CEP123 ortholog, *c14h10.2*, was found to be a novel gene associated with a dye filling defective phenotype (Dyf) and the resistance of C14h10.2(*tm10737*) mutants to avermectins and albendazole support it having a role in the cilia and suggest that it might directly interact with the BBSome or one of the IFT complexes. As T04f8.6(*tm4830*) mutants were susceptible to albendazole it can be deduced that resistance seen in Dnc-1(*or404*) is being caused by a function of the protein unrelated to cilia and is probably fusing using the AEX-4 tSNARE and VAMP-7 vSNARE proteins.

### Transition fibres and the TOGARAM1 complex

The transition fibres attach at the basal body between the axoneme and the ciliary membrane. Through the interaction with nucleoporins (93, 110) and highly redundant interactions among transition fibre proteins (111–113), the transition zone plays a key role in maintaining the impermeability of the ciliary gate. Mutations in transition fibre genes are associated with multiple ciliopathies in vertebrates (112) however, it was surprising that of the transition fibre genes tested only Nphp-4(*tm925*) mutants showed resistance to the avermectins. These findings are consistent with observations of dye-filling defects in *C. elegans* transition fibre mutants, where the high degree of redundancy requires the loss of multiple transition fibre proteins to cause any significant ciliary defects (114). Both the transition fibres and basal body interact with a protein complex involved in the post-translational modification of axoneme tubulin, with CHE-12 being an ortholog of the TOGARAM1 subunit (115). When this complex was investigated only the ARMC9/JBTS-30 ortholog *f59g1.4* was found to cause albendazole resistance, and none of the tested subunit mutants were resistant to ivermectin/moxidectin as observed in Che-12(*e1812*). This suggests highly specialised roles for the core subunits in ciliary maintenance while the periphery subunits exhibit redundancy between the basal body and transition fibre interactions.

There are several transition fibre associated proteins whose specific protein-protein interactions with the basal body have yet to be determined, however our results identified four proteins which have roles in anthelmintic resistance. OSTA-1 is known to cause minor ciliary distal segment length defects, especially in AWB neurons, through RAB-5 mediated regulation of IFT (116). This reduction in distal segment surface area in the OSTA-1 mutant would lead to lower moxidectin uptake and hence may explain the low level of moxidectin resistance. The transition fibre associated protein GASR-8 is orthologous to proteins that form ciliary nexin links through microtubule interaction and bundling in other organisms (106, 117) and it was therefore surprising to find strong levamisole but not albendazole resistance in the Gasr-8(*gk1232*) mutant suggesting that this protein has additional functions in *C. elegans*. The cause of levamisole resistance seen in T06g6.3(*gk546*) mutants remains elusive since little is known about this protein other than it is enriched in the cilia (35) and interacts with the following proteins: AFD-1, GEI-4, LET-413, LIN-15A, LIN-37, NHR-11 and VAB-3 (118, 119). The transition fibre protein DYF-17 has an as-yet undefined role in distal segment assembly (51) however, orthologs are known to interact with BBS-4 and the axon guidance protein UNC-76 (120) suggesting that it may function through facilitating the gating of the BBSome.

### Exosomes, recycling and degradation pathways

In neurons, proteins are synthesised in the soma and require kinesins for anterograde transport along the axon to their destination, whereas retrograde transport is carried out by the dyneins (121, 122). The direction of transport is determined by the polarity of the axonal microtubules and is dependent on UNC-33 and UNC-44 (123). This polarity requirement provides an explanation for the observed resistance to ivermectin in Unc-33(*e1193*) and Unc-44(*e1197*) mutants, as the delivery of amphid cilia proteins and the downstream effectors for ivermectin susceptibility would become disorganised in these mutants. In an attempt to identify which kinesins are responsible for delivering these proteins to the end of the axon, the axonal kinesins KLC-1, KLP-6 and UNC-104 (121, 124, 125) were investigated but found not to influence ivermectin tolerance, indicating either functional redundancy or that ciliary proteins are transported by other cytoplasmic kinesin family members. Unlike the other kinesins tested, the kinesin-13 family, of which KLP-7 is a member, has roles in microtubule depolymerisation and primary ciliogenesis (126). As Klp-7(*tm7884*) mutants showed no ivermectin resistance or dye-filling defects it can be concluded that it plays a negligible role in ciliogenesis in *C. elegans* and the observed albendazole resistance is probably being caused by reduced microtubule/free tubulin cycling resulting in a reduction in available unhindered albendazole binding sites.

Similarly, protein degradation also requires the retrieval of damaged and superfluous proteins across the axon from where they are localized. The results indicated that the downstream effectors for ivermectin susceptibility needs to be returned from the cilia for ivermectin efficacy. The GTPase RAB-35 is involved in the recycling of endosomes which can contain ciliary membrane proteins (127), and resistance to avermectins in this mutant may infer an important role for endocytosis in the efficacy of this class of anthelmintics. Several proteins commonly involved in membrane protein endocytosis were also investigated, however both Cav-1(*ok2089*) and Dpy-23(*e840*) (an AP-2 subunit) mutants remained susceptible to the tested anthelmintics. Surprisingly Chc-1(*b1025*) mutants, for the clathrin heavy chain, were likewise susceptible to all anthelmintics tested. CHC-1 plays an important role in UNC-101, DPY-23 and CAV-1 mediated vesicle formation therefore indicating that vesicle formation for IFT and the downstream effector for ivermectin susceptibility may be occurring via one or more of the clathrin-independent pathways (128).

The Rab GTPases, of which C. elegans has 31 members (129), play important roles in regulating vesicle trafficking and membrane fusion. As RAB-35 was found to have roles in anthelmintic resistance, a selection of key Rabs which function upstream and downstream were investigated to clarify which endosome recycling routes were important for each class of anthelmintic. RAB-2 (also known as UNC-108) is involved in deciding if late endosomes are sent for degradation in the lysosome (130) and RAB-11.1 and RAB-11.2 which facilitate the slow endosome recycling pathway (127) had no impact on anthelmintic resistance indicating that the observed resistances are being caused by defects in the fast recycling pathway. Interestingly Rab-3(*y250*) which regulates the exocytosis of secretory vesicles (131) showed resistance to albendazole despite being a neuron specific RAB in *C. elegans* (132) suggesting that the Unc phenotype observed during exposure in wild types is being caused by microtubule disruption in the nervous system rather than the muscles.

Ciliated neurons in *C. elegans* release protein and RNA containing secreted vesicles called exosomes (also known as ectosomes or extracellular vesicles) (133). Exosomes have a role in inter-organism signalling (134, 135) that can cause resistance by becoming decoys, with proteins being used to take up or bind toxic compounds and pathogens into discarded vesicles (136). The endocytosis of released exosomes can also potentially increase susceptibility by increasing the surface area available for uptake. The impairment of IFT or lysosomal degradation pathways are known to stimulate the release of exosomes that contain cilia proteins (137) and ivermectin resistant Rab-28(*ok3424*) mutants are known to have impaired exosome release (68). Components of the *C. elegans* exosome release/uptake pathway were therefore investigated for their roles in anthelmintic resistance. Exosomes are exported from the cilia in a KLP-6 and CIL-7 dependent manner (133), so if exosomes are important for resistance, mutants would either be more resistant due to a reduction in environmental exosomes or would be more susceptible due to the uptake of accumulated exosomes in the amphids. The Lamp1 ortholog *lmp-1* plays a role in exosome uptake (138) as well as being important for lysosome formation and fusion between endosomes and autophagosomes (139, 140). The Cil-7(*tm5848*) and Klp-6(*tm8587*) mutants were found not to be resistant to the avermectins or albendazole and did not have a mortality rate which was discernible from the other susceptible strains tested. The Lmp-1(*ok3228*) and Rab-2(*e713*) mutants were also susceptible to all anthelmintics tested indicating that neither exosomes or the lysosomal degradation pathway are important for anthelmintic uptake. As Cil-7(*tm5848*) mutants were resistant to levamisole it can be deduced that the protein may have additional functions outwith exosome export.

### Genes that cause broad spectrum cross resistance

In this study, three genes were identified (*daf-6*, *inx-19* and *rab-35*) that when mutated caused broad spectrum cross resistance to the avermectins, albendazole and levamisole. The protein DAF-6 plays an important role in lumen formation and the morphogenesis of the amphidial sheath but is also present in other tubular lumens, such as the intestine, which would also potentially be exposed to anthelmintics (141, 142). DAF-6 is predicted to function by inhibiting endocytosis of the extracellular matrix (142), a determining factor in apical-basal polarity of the lumen (143). Although there is strong evidence that the resistance to the avermectins is being caused by amphidial sheath defects, the resistance towards the other anthelmintics seen in Daf-6(e1377) mutants is probably being caused by the reduction in polarity, thus resulting in downstream effectors for anthelmintic uptake being mislocalised on membrane surfaces. The reduced fluorescence observed when exposed to FBI and BABZ compared to the N2 wild type further supports a reduction in uptake as the cause of resistance. The Rab GTPase RAB-35 determines whether to send early endosomes for recycling as opposed to the lysosomal degradation pathway and plays an important role in maintaining membrane receptor populations (127, 144). This function alone could cause the cross resistance noted by reducing the number of downstream effector proteins available for anthelmintic uptake or restricting the number of primary targets. However, RAB-35 also plays roles in cell migration, neurite outgrowth and cell polarity (143, 145), all of which could reduce target access or uptake for anthelmintics. It was surprising that Rab-35(*b1013*) showed FBI uptake comparable to the wild type while having greatly impaired BABZ uptake (Fig 2J) suggesting that the observed resistances are being caused by more than just one mechanism. The innexins form intercellular channels that function as gap junctions in neurotransmission, with members such as *unc-7* and *unc-9* being involved in ivermectin resistance through what is believed to be a reduction in the transmission of erroneous excitations caused by neurotoxic anthelmintics (12). This was reflected in their uptake of DiI and FBI which was comparable to the wild type. INX-19 is however functioning through a different mechanism since mutants exhibited dye-filling defects and impaired FBI uptake which would indicate structural abnormalities of the ciliated amphid neurons. The INX-19 gap junctions allow the passage of nucleotide signalling molecules and other small compounds between cells (146, 147) potentially facilitating the neural distribution of lipophilic dyes and anthelmintics. There is also a role that INX-19 plays in determining neural cell fate (147, 148) which could be important for the differentiation into cells involved in dye and anthelmintic uptake. As Inx-19(*ky634*) showed greatly reduced BABZ uptake in the gut there is indication that INX-19 might also have a role outside the nervous system. If the above three genes maintain the same roles in parasitic nematode species of economic or medical importance, then it would be possible for a single mutation to render three of the most widely used anthelmintic families ineffective.

### BODIPY labelled anthelmintic analogs

BODIPY labelled probes have been shown to be biocompatible and successfully applied to a variety of biologically relevant compounds (149) and the results show that they are also applicable to anthelmintics. The use of a fluorescently labelled ivermectin probe allowed the hypothesis that the amphids are the tissue responsible for ivermectin uptake to be visually confirmed while the labelled albendazole probe has shown that albendazole enters via the gut. Differences in the intensity of absorbed probe fluorescence compared to wild type also corresponded well with observed resistance in the tested mutant strains. As the intensity of fluorescence is proportional to concentration at the steady state the question remains open as to whether a reduction in uptake or increase in efflux is responsible for the reduced intensity observed in many of the tested mutants, however, given the functions of the mutated genes it is probably being caused by reduced uptake.

The reduced primary toxicity observed in the FBI probe was to be expected as even small structural changes to the 4’’ position can have large effects on the potency of avermectins (150) and the BODIPY fluorophore is a comparatively bulky chemical group. Given the low solubility of benzimidazoles in aqueous/DMSO emulsions (14) relative to their EC_50_s in susceptible strains it is challenging to assay resistant strains and labelled compounds with reduced toxicity as precipitation occurs before the majority of physiologically relevant doses. This means that at higher doses additional routes of exposure occur through ingestion of and direct contact with the precipitate and that identified EC_50_s are not always practically attainable. As the effectors for ivermectin and albendazole uptake are still unknown it was not possible to design and use self-quenching probes and the BODIPY fluorophore, despite having high fluorescence efficiency, is prone to photodegradation at the meso carbon in the presence of environmental oxygen when in polar (eg. aqueous) solutions (149) meaning that videoing the progressive uptake within individuals was not feasible. Still the approach shows promise for identifying the routes and tissues involved in anthelmintic uptake and testing BODIPY labelled analogs of other classes of anthelmintics will be the subject of future work.

### Whole genome sequencing of forward genetics screen mutants

The avermectin resistance causing genes uncovered by random mutagenic screens and identified by whole genome sequencing were all determined to be involved in IFT. The dynein heavy chains *che-3* and *dhc-3* were found to be mutated more commonly than other genes. This is similar to TP238(*ka32*) and TP239(*ka33*) from the previous study looking for ivermectin resistance (23) and other studies looking for dye-filling defects (49, 151). This overrepresentation of dyneins in forward genetic screens is probably caused by dynein heavy chains having very long coding sequences (12,516nt and 9,828nt for *che-3* and *dhc-3* respectively) making them more prone to mutation by EMS as the rate of mutation for a loss of function mutation is proportional to gene size (152). Having found genes involved in ciliogenesis and IFT is not surprising as it is a complex process requiring the interaction of multiple genes to produce a functional structure and lacks redundancy. Therefore, for the aforementioned reasons, genes involved in IFT are statistically more likely to be mutated by EMS than single downstream effectors that rely on functional cilia.

## Conclusion

The findings of this study not only provide strong evidence that the avermectin compounds ivermectin and moxidectin are taken up via the amphid cilia as has been shown previously (23) but refines the location of the effectors to the distal segment of the cilia. This study also uncovers the pathways used to deliver the effectors and other ciliary proteins in *C. elegans*. Due to the strong correlation between IFT function with dye-filling defects and resistance to avermectins it may be possible to use resistance phenotypes to identify if novel dye-filling mutants from forward genetic screens are upstream or downstream of IFT. There is also evidence that the three complexes associated with IFT have additional roles in protein trafficking outside of IFT and these may be responsible for resistance to albendazole. Levamisole was not the primary focus of this study however several genes associated with ciliary processes were found to cause resistance or increased susceptibility indicating a sharing of proteins by other pathways, including those needed to compensate for the loss of controlled cholinergic neurotransmission. Although the downstream effectors for ivermectin uptake remains elusive it can be deduced from the chemical properties of ivermectin (153) that such effectors must associate with the extracellular membrane. The results from this study suggest that the effectors localise to the distal segment of the amphid cilia and possess either a transmembrane domain or are anchored via a myristoyl or palmitoyl group. Whether the effectors are functioning as a carrier protein or transporter remains to be determined. If the resistance causing genes uncovered in this study have the same functions in other nematode species, then there would be important implications for resistance monitoring strategies.

## Acknowledgements

Some strains were provided by the CGC, which is funded by NIH Office of Research Infrastructure Programs (P40 OD010440). Some *C. elegans* strains used in this work were created by the International *C. elegans* Gene Knockout Consortium. Some strains were provided by the National BioResource Project (Japan). Many thanks to Eric Jorgessen for Dyf-17(*ox175*) and Jinghua Hu for Dyf-19(*jhu455*).

---

**Fig S1. Timecourse of BODIPY labelled ivermectin analog, FBI, probe localisation and uptake.**

Timecourse of FBI uptake in N2 at (A) 6 hours, (B) 12 hours, (C) 18 hours, (D) 24 hours, (E) 30 hours, (F) 36 hours, (G) 42 hours, (H) 48 hours, (I) 54 hours, (J) 60 hours, (K) 66 hours and (L) 72 hours. Individuals were photographed using a DIC filter (lower right inset image) to highlight the position and orientation of the worm and a FITC filter (main image) to visualise fluorescence. Areas of weak fluorescence are highlighted with arrows.

**Fig S2. Timecourse of BODIPY labelled albendazole analog, BABZ, probe localisation and uptake.**

Timecourse of BABZ uptake in N2 at (A) 100x magnification at 1 hours, (B) 250x magnification of gut at 1 hours, (C) 100x magnification showing transient uptake at 1 hours, (D) 100x magnification at 2 hours, (E) 250x magnification of gut at 2 hours, (F) 100x magnification at 3 hours, (G) 250x magnification of gut at 3 hours. Arrow highlights observed uptake boundary, (H) 250x magnification of eggs in the body cavity at 3 hours, (I) 100x magnification at 4 hours, (J) 250x magnification of gut and eggs in the body cavity at 4 hours, (K) 100x magnification at 5 hours, (L) 250x magnification of eggs in the body cavity at 5 hours, (M) 100x magnification at 6 hours (N) 250x magnification of gut at 6 hours (O) 250x magnification of eggs in the body cavity at 6 hours, (P) 100x magnification at 7 hours, (Q) 250x magnification of head and gut at 7 hours, (R) 100x magnification at 8 hours, (S) 250x magnification of head, gut and eggs in the body cavity at 8 hours. Individuals were photographed using a DIC filter (lower right inset image) to highlight the position and orientation of the worm and a FITC filter (main image) to visualise fluorescence.

**Fig S3. Schemes for BODIPY labelled anthelmintic analog probe synthesis.**

Schemes for FBI synthesis (A) and BABZ synthesis (B).

